# *TaEXPB5* is responsible for male fertility in thermo-sensitive male-sterility wheat with *Aegilops kotschyi* cytoplasm

**DOI:** 10.1101/2021.12.24.474147

**Authors:** Xingxia Geng, Xiaoxia Wang, Jingchen Wang, Xuetong Yang, Lingli Zhang, Xiyue Song

## Abstract

Thermo-sensitive male sterility is of vital importance to heterosis, or hybrid vigor in crop production and hybrid breeding. Therefore, it is meaningful to study the function of the genes related to pollen development and male sterility, which is still not fully understand currently. Here, we conducted comparative analyses to screen fertility related genes using RNA-seq, iTRAQ, and PRM-based assay. A gene encoding expansin protein in wheat, *TaEXPB5*, was isolated in KTM3315A, which was in the cell wall and preferentially upregulated expression in the fertility anthers. The silencing of *TaEXPB5* displayed pollen abortion, the declination or sterility of fertility. Further, cytological investigation indicated that the silencing of *TaEXPB5* induced the early degradation of tapetum and abnormal development of pollen wall. These results revealed that the silencing of *TaEXPB5* could eliminate the effects of temperature on male fertility, and resulting in functional loss of fertility conversion, which implied that *TaEXPB5* may be essential for anther or pollen development and male fertility of KTM3315A. These findings provide a novel insight into molecular mechanism of fertility conversion for thermo-sensitive cytoplasmic male-sterility wheat, and contribute to the molecular breeding of hybrid wheat in the future.

**Highlight:** *TaEXPB5* coffers to anther or pollen development and male fertility in KTM3315A, its silencing could eliminate the effects of temperature on male fertility, and resulting in functional loss of fertility conversion.

## Introduction

Utilization of heterosis is an effective way to improve food production (Kratochvil *et al*., 1990). Hybrid seed production by cytoplasmic male sterile lines is an important aspect of crop heterosis utilization, and it also has important applications and practical value in hybrid breeding of crop, especially in rice, maize, and wheat (Li *et al*., 2016).

The main pathways for the utilization of heterosis in wheat currently include ‘three-line’, ‘two-line’ and chemical hybridization (Sang *et al*., 2005). To date, ‘three-line’ system has some shortcomings, such as the complicated breeding procedure and high cost of seed production (Guan *et al*., 2002). For chemical hybridization pathway, because most of the chemical pesticides tested are not completely effective, or there is no safety guarantee for humans and animals, resulting in the popularization and application of chemical pesticides is still limited (Jiang *et al*., 2009). However, hybrid breeding based on two-line pathway not only simplifies the breeding program of the sterile line, but also ensures seed purity of the line, therefore, which is the most widely used and most effective procedure.

KTM3315A, a thermo-sensitive cytoplasmic male sterile line with *Aegilops kotschyi* cytoplasm, was developed using TM3315B as a ponor parent for continuous backcrossing. The thermo-sensitivity is determined by *rfv*_*1*_^ma^, which comes from the short arm of 1B chromosome in *Triticum macha* (Song *et al*., 2013). Our previous studies showed that the fertility of KTM3315A is sensitive on temperature during Zadoks growth stages 45 to 52, and it exhibited complete sterility when the temperature was below 18°C, but which has the capacity to self-pollinate when the temperature is above 20 °C (Meng et al., 2016). Therefore, the ‘two-line’ hybrid system based on the thermo-sensitive cytoplasmic male sterility has important application potential in hybrid wheat breeding. At present, there were many studies on male sterility, but mainly focused on their phenotype, physiology, cytology and mRNA expression levels (Meng *et al*., 2016; Ye *et al*., 2017). Some studies on the mechanism of male sterility at the protein level have also been carried out, mainly using traditional 2D gel electrophoresis (2-DE) (Wang *et al*., 2015). This method has many limitations in protein identification, such as difficulty in identifying low-abundance proteins, low protein resolution, and complex protein separation steps (Molloy *et al*., 1998; Ge *et al*., 2013). Nevertheless, these studies provide a reference for the study of male sterility mechanisms.

With the development of high-throughput sequencing technology, proteomics is widely used in fertility studies of a variety of plant. Some fertility-related proteins have also been identified, which are involved in different metabolic pathways, including carbohydrate metabolism and energy metabolism (ATP synthesis subunits) in wheat (Zheng *et al*., 2014; Geng *et al*., 2018), flavonoid synthesis in CMS *Brassica napus* (Sheoran *et al*.,2010) and watermelon (Rhee *et al*., 2015), and transcription factor subunits in *Brassica napus* (Sheoran *et al*., 2010). Isobaric tags for relative and absolute quantification (iTRAQ) is one of the most effective methods for identifying protein profiles, which can be used for the analysis and identification of differential proteins in different tissues or cells. Many studies on the mechanism of male sterility have been carried out using iTRAQ technology, which has been quite successful in discovering new candidate genes and generating hypotheses for fertility conversion. Despite numerous reports describing the fast-growing application of iTRAQ-based quantification of target proteins in crops (Zheng *et al*., 2014; Ji *et al*., 2015; Li et al., 2016), the research on crop male fertility is very limited and has not been applied to fertility conversion in wheat. Parallel reaction monitoring (PRM), a quadrupole-equipped high resolution and accurate mass instruments, was propose as a promising new addition to the quantitative proteomics toolbox for quantifying relative abundance level of proteins due to PRM’s high selectivity. It can precisely quantify selected proteins in complex cellular backgrounds with absolute specificity and sensitivity using full and mass range scans as survey scans together with data-dependent (DDA) and targeted MS/MS acquisition (Amelia *et al*., 2012; Majovsky et al., 2014; Chen *et al*., 2017; Bostanci *et al*., 2018).

With the advent of functional genomics, many predicted gene functions are waiting to be revealed. However, gene function research methods such as antisense RNA technology, gene overexpression, gene knockout, or mutant library were more and more ineffective in identifying the gene function of wheat. Virus-induced gene silencing (VIGS) has the advantages of simple, fast, effective and high throughput because it does not acquire genetic transformation or mutant plants. It has been applied more and more widely in the study of plant gene function (Senthil-Kumar and Mysore 2014). VIGS technology refers to the use of recombinant viruses carrying foreign target gene fragments to infect plants and inhibit the expression of genes that are homologous or identical to the inserted fragment sequence (Burch-Smith *et al*., 2004; Robertson 2004). Barley stripe mosaic virus (BSMV) is a short rod RNA virus, whose genome is composed of three RNA chains, α, β and Y. In 2002, BSMV vector for VIGS was successfully constructed for the first time, and BSMV-VIGS gene silencing system mediated by BSMV was successfully applied to monocotyledons of barley. Subsequently, BSMV-mediated gene silencing was found in other monocotyledons of *Hordeum vulgare* (Scofield *et al*., 2005). *Brachypodium distachyon* (Pacak *et al*., 2010), *Triticum dicoccoides* (Scofield and Nelson 2009), *Haynaldia villosa* (Wang *et al*., 2010) and *Lolium perenne* (Martin *et al*., 2013). In addition, the BSMV-VIGS system in roots of wheat and oat was firstly established, and verified the silencing effect of BSMV-VIGS system in roots by inhibiting the expression of *PHR1* and *PH02* genes (Pacak *et al*., 2010) Furthermore, an effective virus-induced silencing system of endogenous gene in wheat spikes was established, and used this method to silence the glutenin genes *HMW-1Bx14* and X-type HMW-GS gene family, which are known to function in wheat grains, and confirmed that this method can be effectively applied to the functional study of wheat grain genes (Ma *et al*., 2012).

In this study, we performed transcriptome and proteome sequencing of KTM3315A under different fertility conditions using RNA-Seq and iTRAQ techniques, and identified some candidate genes related to fertility conversion. Combined with qRT-PCR assay and PRM-based protein relative abundance analysis, a gene involved in anther development and male fertility, *TaEXPB5*, was validated, and its potential cytological mechanism of anther or pollen abortion was revealed. This study will provide a novel insight into the fertility mechanism of thermo-sensitive cytoplasmic male sterility in wheat.

## Materials and Methods

### Plant material and growth conditions

In this experiment, KTM3315A, a thermo-sensitive male sterile wheat line, TM3315B, the maintainer line of KTM3315A, BSMV:0, an infected plant by BSMV:α/β/γ under fertile condition, and as well as BSMV:*TaEXPB5*, an infected plant by BSMV:α/β/γ-*TaEXPB5* under fertile condition, were used as materials. In early October 2019, KTM3315A were planted in pots and fields at the experimental fields at Northwest A&F University in Yangling (34°29 ‘ N, 108°’ 08E). On April 2, 2020, all the flower pots were moved into two climate incubators, 15 pots per incubator, where day and night time was set to 14 hours and 10 hours respectively, and the corresponding temperature was set to 17°C/15°C for sterile conditions (AS) and 22 °C/20 °C for fertile conditions (AF). The anthers of silencing plants (BSMV:*TaEXPB5*) and control plants (AF and BSMV:0) at the five developmental stages (Tds, tetrad stage; Euns, early uninucleate stage; Luns, later uninucleate stage; Bns, binucleate stage; and Tns, trinucleate stage) were placed in centrifuge tubes for storage in FAA fixative and 4% glutaraldehyde for subsequent cytological analysis, in which the pollen developmental stages were described previously (Song *et al*., 2014). The anthers of AS, AF, and TM3315B in Euns, Bns, and Tns were applied to RNA sequencing and iTRAQ analysis to identify candidate genes related to anther development and male fertility in KTM3315A under different fertility conditions.

### Anther phenotyping

Anthers were observed and photographed using a Motic K400 dissecting microscope (Preiser Scientific, Louisville, KY, USA), which equipped with a Nikon E995 digital camera. Fresh microspores were stained with iodine-potassium iodide (2% I2-KI) to detect pollen granule activity. The anthers and microspores in Tns were observed with a JSM-6360LV scanning electron microscope (SEM) and photographed. The treatment of anthers and microspores was basically as described (Zhang et al. 2010; Xu et al. 2014).

### RNA sequencing, iTRAQ sequencing, and PRM-based quantitative proteomic analysis

Anthers of Euns, Bns, and Tns in KTM3315A under sterile condition (AS1, AS2, and AS3), KTM3315A under fertile condition (AF1, AF2, and AF3), were perfomed RNA sequencing (RNA-seq) as previous descripted (Ye *et al*., 2017). Proteins from anthers of AS (AS1, AS2, and AS3), and AF (AF1, AF2, and AF3), and TM3315B (B1, B2, and B3) were extracted using the method as described (Geng *et al*., 2018). For PRM-based quantitative proteomic analysis, 18 target proteins in 9 samples (A1, A2, A3, AF1, AF2, AF3, B1, B2, and B3) were quantified by PRM. The samples were fine grinded by liquid nitrogen into powder, and added four volumes of lysis buffer (8 M Urea, 1% Triton-100, 10 mM Dithiothreitol, and 1% Protease Inhibitor Cocktail). After following by sonication three times on ice using a high intensity ultrasonic processor (Scientz), add an equal volume of Tris equilibrium phenol, 4 °C, 5500 g, centrifuge for 10 min, collect the supernatant and add 5 times the volume of 0.1 M ammonium acetate or methanol for precipitation overnight, and wash the protein precipitation with methanol and acetone respectively. Finally, the protein was redissolved in 8 M urea and the protein concentration was determined with BCA kit according to the manufacturer’s instructions. For digestion, the protein solution was reduced with 5 mM dithiothreitol for 30 min at 56 °C and alkylated with 11 mM iodoacetamide for 15 min at room temperature in darkness. The protein sample was then diluted to urea concentration less than 2M. Finally, trypsin was added at 1:50 trypsin-to-protein mass ratio for the first digestion overnight and 1:100 trypsin-to-protein mass ratio for a second 4 h-digestion. Then LC-MS analysis was performed, the resulting MS data were processed using Skyline (v.3.6).

### Green fluorescent protein (GFP) subcellular localization

The subcellular localization expression vector was constructed using the CDS of the *TaEXPB5* fragments that were cloned into the *Xba* I/*Xho* I site of the digested pKANNIBAL27 to construct the pKANNIBAL27-*TaEXPB5* vector. Then the constructed *TaEXPB5*-GFP and GFP were transferred to the *Agrobacterium tumefaciens* GV3101. The free GFP was as control. The cultured cells were resuspended in buffer [10 mM MgCl2, 10 mM 2-(N-morpholino) ethanesulfonic acid (MES), pH 5.6, and 0.2 mM acetosyringone (AS)], where the OD600 was adjusted to 0.3, then infiltrated into *N. benthamiana* leaves using a needleless syringe. A confocal laser scanning microscope (IX83-FV1200) under the excitation of 488 nm and emission of 587 nm were used to observe the fluorescence. The primer sequences involved in the experiment were shown in Supplementary Table S1.

### Virus induced gene silencing by barley stripe mosaic virus

Gene silencing assay involved in α, β, and γ vectors of barley stripe mosaic virus (BSMV). Herein, γ-*TaPDS* and γ-*TaEXPB5* recombinant vector were constructed by inserting into the γ vector with the conserved sequence of *TaPDS* and target *TaEXPB5* gene, respectively. In vitro, the three transcripts of α/β/γ mixed with FES buffer at the ratio of 1:1:1:18, as were that of α/β/γ-*PDS* and α/β/γ-*TaEXPB5*. Subsequently, each transcript mixture infected the flag leaves and the first two leaves of wheat at the booting stage, and cultivated at day/night temperatures of 22°C/25°C. The plants infected by the three combinations were named BSMV:0 (α/β/γ), BSMV:*TaPDS* (α/β/γ-*PDS*), BSMV:*TaEXPB5* (α/β/γ-*TaEXPB5*).

### Quantitative RT-PCR analysis of mRNA

The total RNA was extracted using the RNA prep pure plant kit (Tiangen Biotech (Beijing) Co., Ltd.), reverse-transcribed (RT) using a Transcriptor First Stand cDNA synthesis kit (Roche) and quantified on Quantitative PCR Q7 Detection System (Thermo Fisher Scientific, Waltham, USA) with the 2×RealStar Gree Fast Mixture (Genstar (Singapore) Investment Pte. Ltd.). The wheat actin gene was used as the reference gene, in which three independent biological replicates were conducted per analysis with at least three technical replicates. The primers have been listed in Supplementary Table S1.

### Cytological observations

The anthers of silencing plants (BSMV:*TaEXPB5*) and control plants (AF and BSMV:0) at different stages were fixed in FAA solution and then embedded in paraffin for cross-section observation, as described (Wang et al. 2016). The analysis of microspores at trinucleate stage were carried out by staining with 4’,6-diamidino-2-phenylindole (DAPI).

For observing the development of anthers by ultra-thin sections, the anthers were fixed, dehydrated, infiltrated and embedded according to Yang et al. (2018), and stained followed by Wang et al. (2013). Ultrathin section observation and image capture were carried out by a JEM-1230 transmission electron microscope (TEM) (JEOL, Tokyo, Japan).

### Terminal deoxynucleotidyl transferase-mediated dUTP nick end labelling assay

For the Terminal deoxynucleotidyl transferase-mediated dUTP nick end labelling (TUNEL) assay, Paraffin sections were dehydrated, washed, dyed and observed according to the procedure (Liu et al. 2018), and assessed using fluorescence confocal scanner microscope (A1R; Nikon, Tokyo, Japan), in which the emission/excitation spectrum of green fluorescence of fluorescein (TUNEL signal) and blue fluorescence of DAPI were analyzed at 450/515 nm, and at 358/461 nm, respectively (Wang et al., 2015).

### DNA laddering analysis

The DNA fragmentation was evaluated as previously described (liu et al., 2018) with minor modification. Briefly, 10 ug DNA of anthers from the different development stages in positive control (AF and BSMV:0) and the silencing plants (BSMV:*TaEXPB5*) was used for electrophoresis on 1.8% (w/v) agarose gel with 0.5 × TBE electrophoresis buffer, and observed and photographed.

### Statistical analyses

All the assays were carried out at least three times, and the data were displayed as mean ± standard deviation (SD). All statistical analyses were performed by Student’s t-test in SPSS statistical software (IBM, NY, USA). *p* < 0.05 and < 0.01 represented significant differences and extremely significant differences, respectively.

## Results

### Phenotyping of KTM3315A under different fertility conditions

To investigate the effects of temperature on male fertility, including anther or pollen development and fertility determination of KTM3315A, we performed thermo-treated assay for KTM3315A (day/night temperature, 22°C/20°C and 17°C/15°C). The result showed that KTM3315A (AS) treated under 17°C/15°C temperature displayed anthers slender, not dehiscent (Fig. 1A1), incomplete staining pollens with potassium iodide (Fig. 1A3), and two abnormal round sperm nuclei at trinucleate stage (Fig. 1A2), while KTM3315A (AF) treated under 22°C/20°C temperature with hypertrophic and dehiscent anthers (Fig. 1B1), complete staining pollens with potassium iodide (Fig. 1B3), and two normal spindle sperm nuclei (Fig. 1B2). Further, AS showed complete male sterility (Fig. 1A4) and AF displayed fertility restoration (Fig. 1B4). This result confirmed the thermo-sensitivity of TCMS-K line KTM3315A, which can occur fertility conversion events and have fertility phenotype at higher temperatures.

**Fig. 1.**
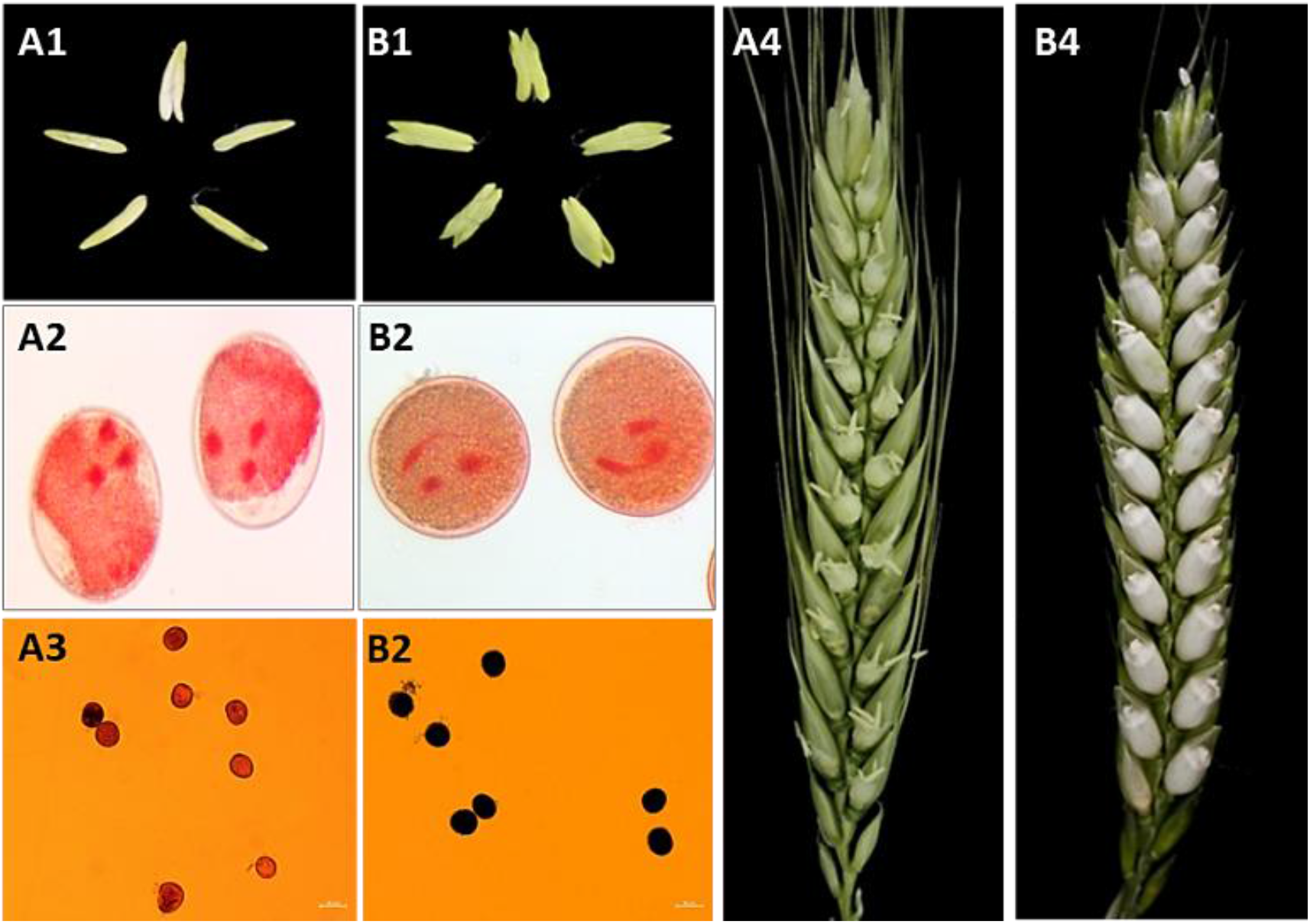
Anther observation and pollen vitality detection in KTM3315A under different fertility conditions. (A1-A4) AS, KTM3315A under sterile conditions; (B1-B4) AF, KTM3315A under fertile conditions; (A1, B1) anthers at trinucleate stage, the anthers of the AS were not cracked, the anthers from AF were dehiscent with shedding of the mature pollen grains. (A2, B2) pollens using magenta acetate stains, the sperm nuclei were round in the AS instead of being fusiform in shape. (A3, B3) pollens stained by potassium iodide, AS pollen could not be stained deeply, AF pollen was rounded with full and deep dyeing. (A4, B4) Seed setting rate, the seed setting rate of AS was 0%, the seed setting rate of AF was 98%. The scale bars indicated are 50mm (A1-A3, B1-B3).

### TaEXPB5 was upregulated in the anthers of KTM3315A under fertile condition, and significantly abounded for its encoded protein

To obtain genes related to anther development and fertility conversion, we performed RNA sequencing for AS and AF, and iTRAQ analysis for AS, AF, and B respectively, where FDR (False Discovery Rate) ≤ 0.05 and |Fold Change| ≥ 2.0 (≥ 1.2 for iTRAQ) were used as thresholds for detecting differential expression genes. According to RNA-seq and iTRAQ analysis, *TaEXPB5*, a gene encoding expansin protein in wheat, was identified, and upregulated expression in fertility anther of AF under fertile condition (Fig. 2A). Also, PRM-based assay (Mass spectrometry based targeted proteome quantification) confirmed that the TaEXPB5 significantly abounded in fertility anthers of Bns and Tns (Fig. 2B), indicating that *TaEXPB5* may involve in regulating male fertility of KTM3315A against different fertility conditions.

**Fig. 2.**
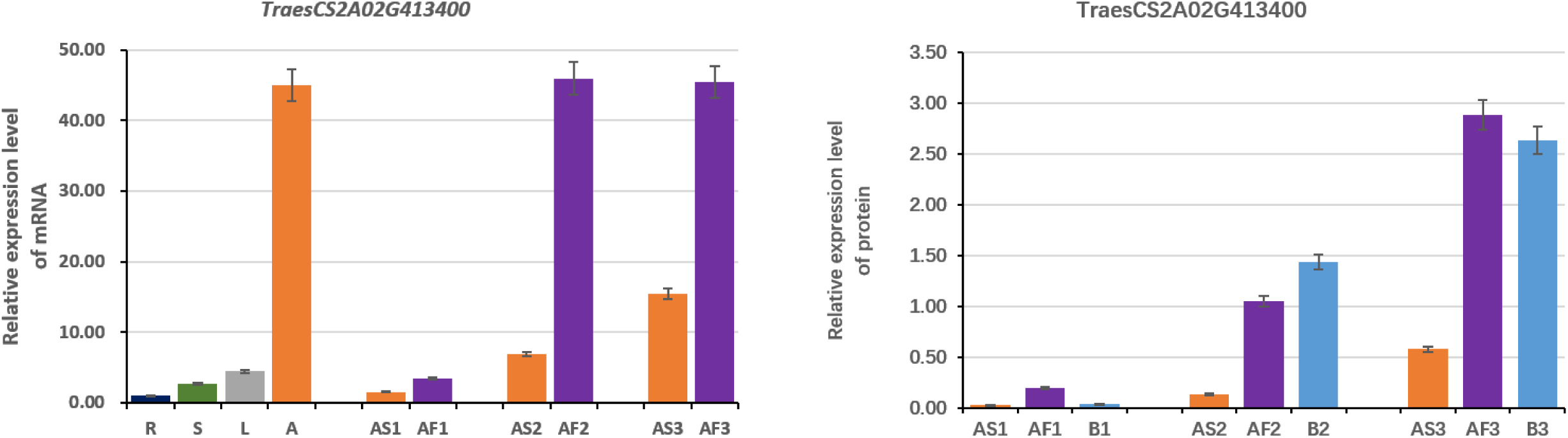
The spatiotemporal expression levels and protein-relative abundance level of TaEXPB5 (TraesCS2A02G413400) KTM3315A at earlier uninucleate stage (Euns), binucleate stage (Bns), and trinucleate stage (Tns) under different fertility conditions. AS1, AS2, and AS3 presents KTM3315A at Euns, Bns, and Tns under sterile condition, respectively; AF1, AF2, and AF3 separately refer to KTM3315A at Euns, Bns, and Tns under fertile condition; B1, B2, and B3 separately refer to TM3315B at Euns, Bns, and Tns. R, root. S, stem. L, leaf. A, anther.

### TaEXPB5 conferred to pollen development and male fertility under fertile condition

To further confirm the candidate gene, the full-length coding sequences of *TaEXPB5* were cloned from KTM3315A under different fertility conditions (AS and AF). The sequence analysis showed that CDS (786 bp) of *TaPEXB5* in AS is the same as that in AF, encoding 261 amino acids (Fig. 3A; Supplementary S1). The predicted protein contains a conserved DPBB_EXPB_N (Accession<u>:cd22275</u>) domain and it belongs to the DPBB_RlpA_EXP_N-like superfamily (Fig. 3A), *TaEXPB5* may be involved in cell wall formation. To further understand the evolutionary relationship between TaEXPB5 and other species homologous genes, and predict its potential function, through blast search against the WheatOmics 1.0 (http://202.194.139.32/), a total of 11 homologous genes were obtained. The neighbor-joining (NJ) phylogenetic trees were constructed by MEGA 7.0 (Fig. 3B). the result revealed it shares a close relationship with AT1G65681 (EXPB6), LOC_Os02g44108 (OsEXPB11), and AT2G45110 (EXPB4). Indicating that TaEXPB5 may have same function as them (Fig. 3B). Its position (cell wall and nuclei) was confirmed by subcellular localization assay (Fig. 4), and which was specifically upregulated for male fertility of KTM3315A under fertile condition (Fig. 2). Therefore, suggested that *TaEXPB5* may be related to pollen development and male fertility.

**Fig. 3.**
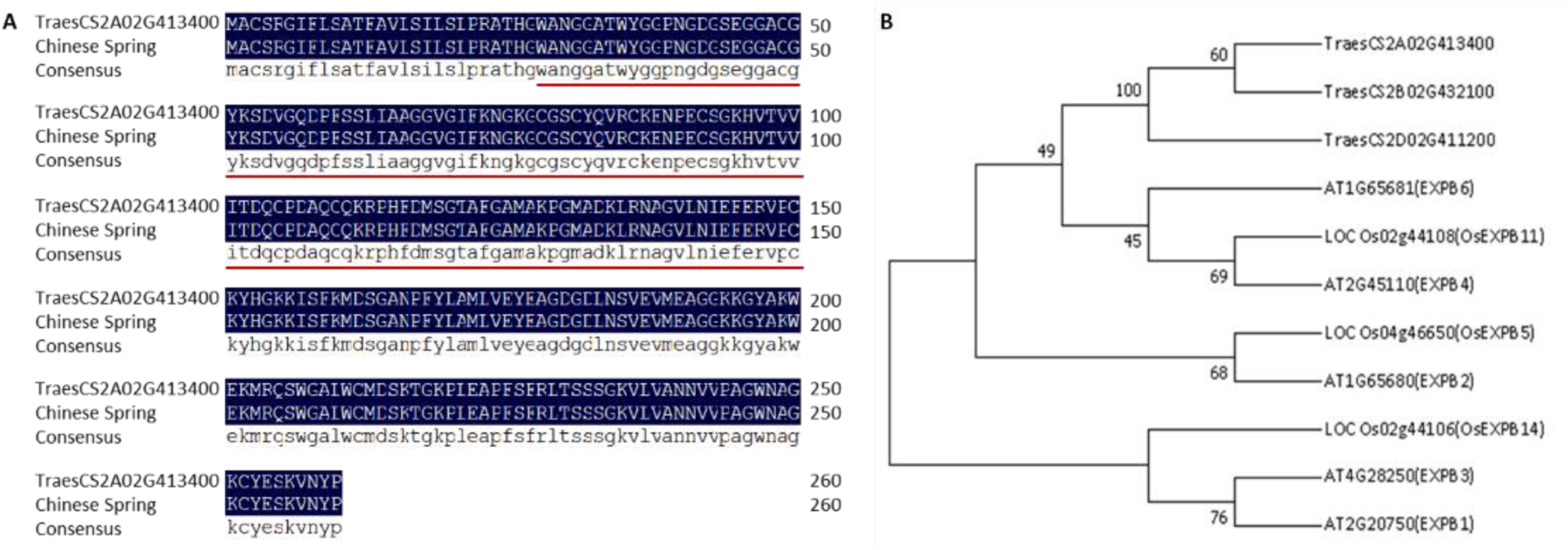
Gene structure and evolutionary tree. (A) Alignment of amino acid sequences for TaEXPB5 in KTM3315A and the reference genome sequence (IWGSC_v1.1_LC_gene). The red line indicates the conserved DPBB_EXPB_N region that belongs to the DPBB_RlpA_EXP_N-like superfamily. (B) Bayesian phylogenetic tree of all of the homologous genes from *Triticum aestivum, Oryza sativa, Arabidopsis thaliana*, it shares a close relationship with AT1G65681 (EXPB6), LOC_Os02g44108 (OsEXPB11), and AT2G45110 (EXPB4).

**Fig. 4.**
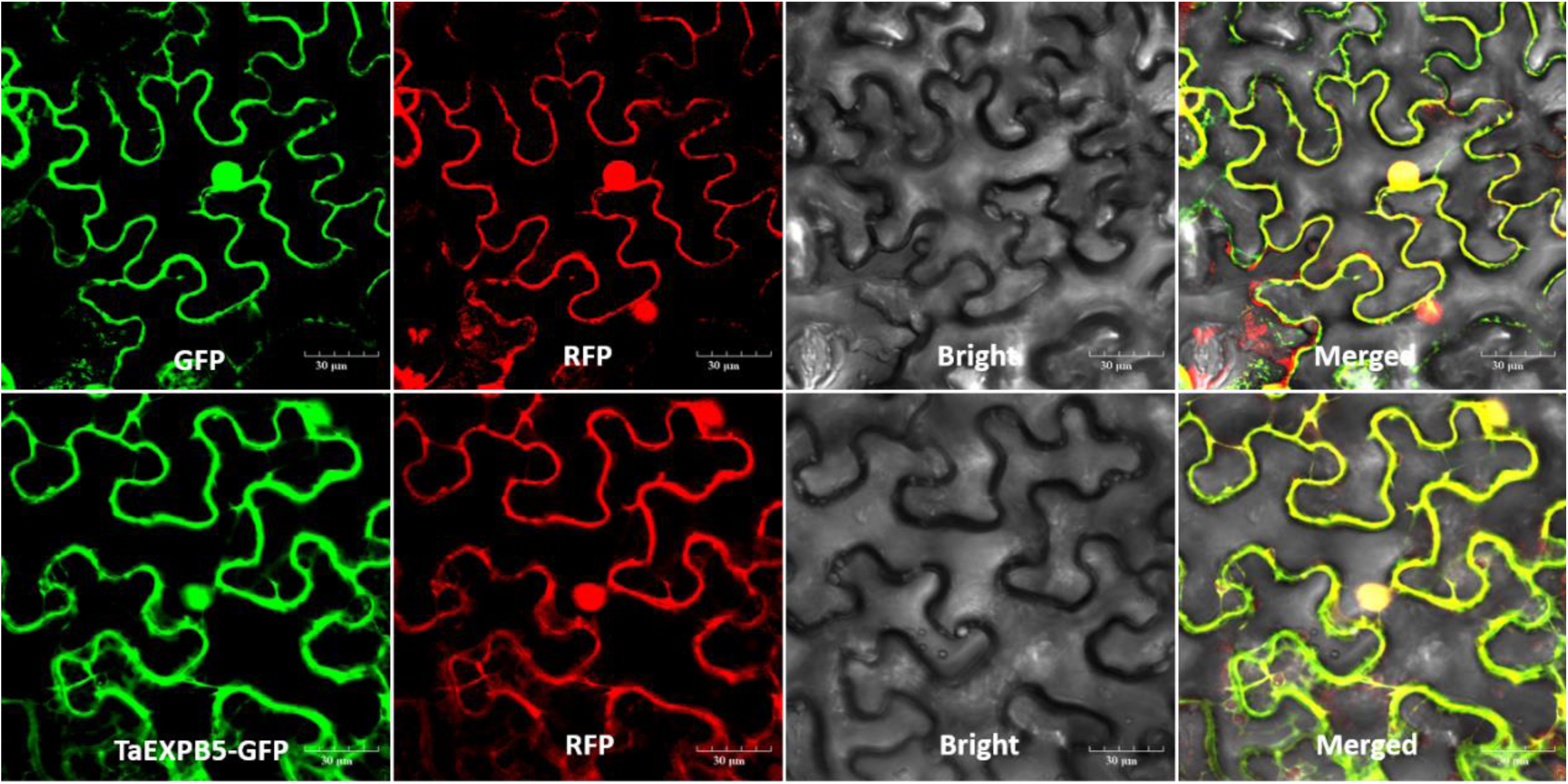
Subcellular localization in epidermal cells of tobacco leaves. (A) Control (35S-GFP), the green fluorescent protein signal filled the whole cell. (B) Co-expression of TaEXPB5-GFP, Plasma membrane-maker (mCherry) and nuclei-marker (HY5), the green fluorescent signal expressed by TaEXPB5-GFP was fused with the red fluorescent signal of Plasma membrane-maker and nuclei-marker. Scale bars are 30µm.

To further confirm whether *TaEXPB5* confer to pollen development and male fertility, we performed gene silencing assay of *TaEXPB5* by BSMV-VIGS (Fig. 5; Supplementary S2). Compared with the control plants, the anthers of silencing plants were smaller and did not crack (Fig. 5c), and self-seed setting was hardly zero (Fig. 5B). The expression reducing of *TaEXPB5* in silencing plants further confirmed its silencing (Supplementary Fig. S3). Indicating that *TaEXPB5* play a role on anther development and male fertility in KTM3315A.

**Fig. 5.**
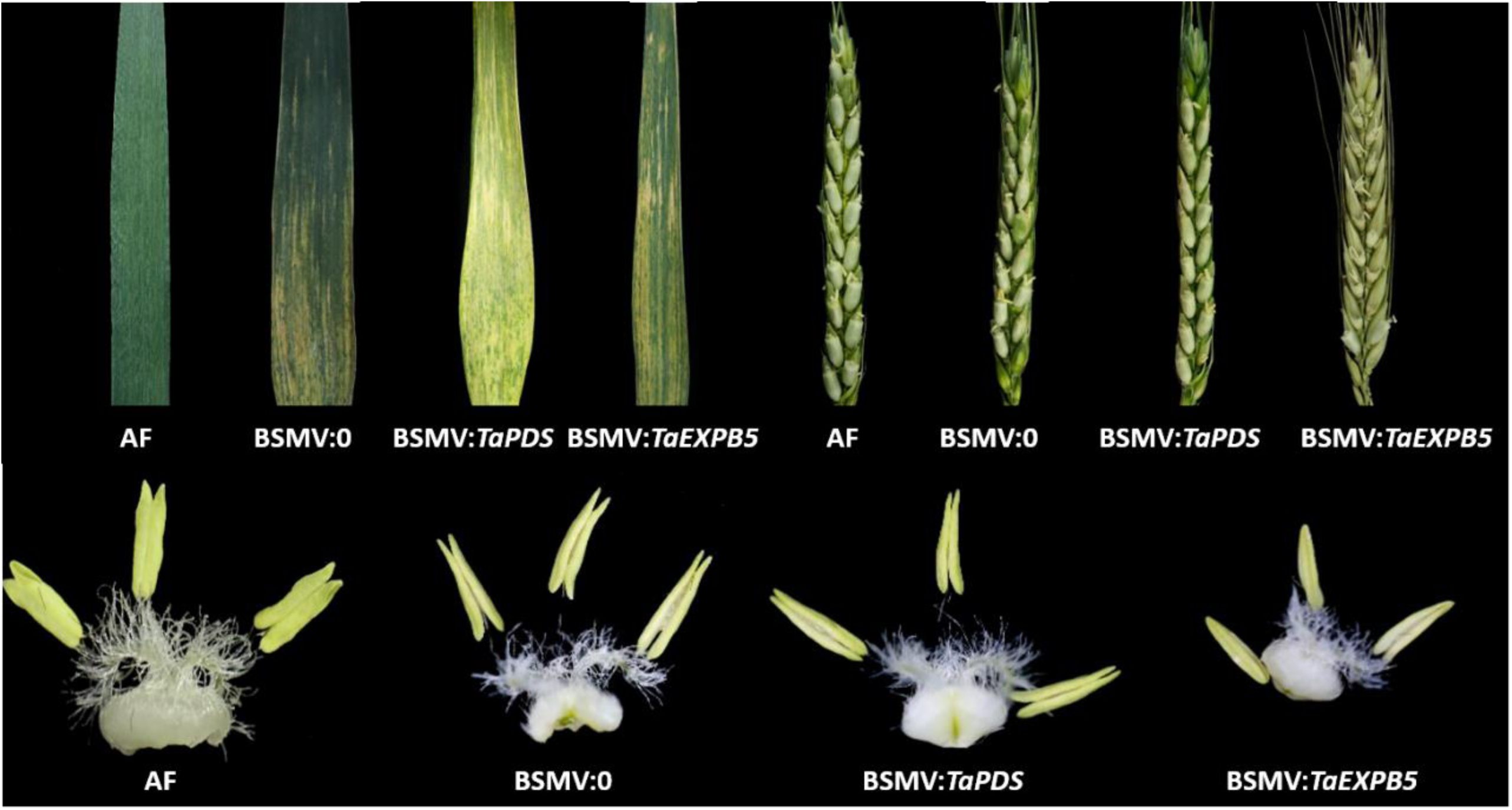
Phenotypic observation of silencing plants BSMV:*TaEXPB5* and control plants (BSMV:0, AF). (A) leaves. It was found that the leaf phenotype of uninfected plants was normal: the leaf color was dark green without any abnormality. The infection of BSMV: 0 resulted in streaks in the leaf. The plants inoculated with BSMV:*TaEXPB5* showed green fading and a yellow stripe appearance. (B) Seed setting rate. Contrary to control plants, the self-seed setting rate of BSMV:*TaEXPB5* was hardly zero. (C) anthers at trinucleate stage. The anthers of the control plants were dehiscent with shedding of the mature pollen grains, and the anthers from BSMV:*TaEXPB5* not cracked.

### TaEXPB5 is required for pollens development and male fertility under fertile condition

Previous studies have shown that genes encoding expansin protein are closely related to plant wall development and fertility (Liu *et al*., 2021). Here, to investigate whether *TaEXPB5* is related to pollen wall development and male fertility as previously described, we conducted the following experiments, including observing the external morphology, outer epidermis, and microspores of Tns anthers by SEM after *TaEXPB5* silencing (Fig. 6). The results showed that the anthers of silencing plants were more shriveled and smaller than that of control plants (Fig. 6A-C), and as well as with the wrinkle epidermal and disorder cells (Fig. 6A1, A2). Compared with control plants having the microspores of the rounded shape and smooth surface (Fig. 6B3, B4, C3, C4), the surface of the microspores was rough and wrinkled (Fig. 6A3, A4). Moreover, pollen grains can be stained lightly and incompletely by I2–KI (Fig. 6A5), while completely and deeply for that in control plants (Fig. 6B5, C5; Supplementary S3). Then, to further determine male fertility of KTM3315A after the silencing of *TaEXPB5*, we performed DAPI staining assay to observe microspores at Tns. It was found that the sperm nuclei of microspores of silencing plants were significantly different from those of control plants. That is, there were two round sperm nuclei rather than two spindle sperm nuclei (Fig. 6E). Indicating that the silencing of *TaEXPB5* can cause abnormal development of anthers or pollen, thereby resulting in the same male sterility as AS (Fig. 1A1-4). Those results further revealed that *TaEXPB5* are necessary for pollens development and male fertility under fertile condition.

**Fig. 6.**
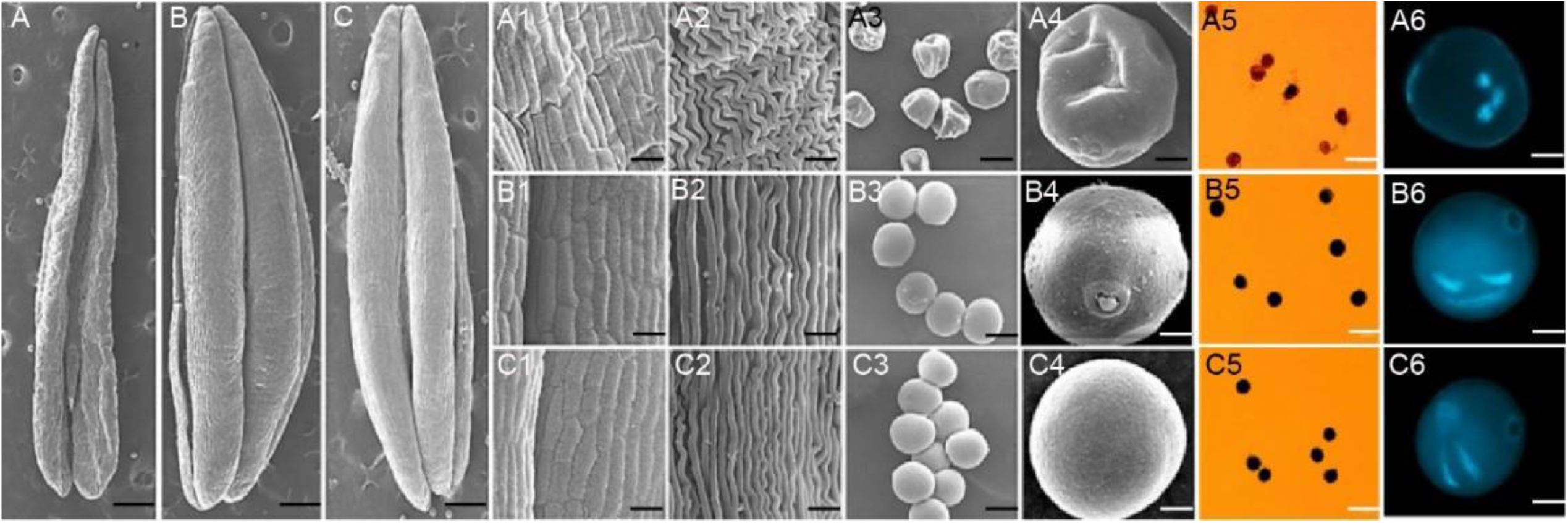
Comparison of scanning electron micrograph observations, I2-KI staining, and DAPI staining in silencing plants and control plants in Tns. (A, A1-A6) silencing plants (BSMV:*TaEXPB5*); (B, B1-B6) control plants (BSMV:0); (C, C1-C6) control plants (AF); (A1, A2, B1, B2, C1, C2) The outer epidermal cells in the anthers were more shrunken than those in control plant cells, and they had irregular shapes. (A3, A4, B3, B4, C3, C4) the epidermis was more rounded and plump with a gymnotremoid germinal aperture in the microspores of control plants, whereas the epidermis was crenate and extremely scabrous with a malformed germinal aperture in the microspores of BSMV:*TaEXPB5*. (A5, B5, C5) In contrast to the pollen of control plant, the pollen grains of BSMV:*TaEXPB5* could not be stained deeply. (A6, B6, C6) The pollen of control plant contained two fusiform sperm nuclei and a vegetative nucleus, whereas the sperm nuclei were round in the BSMV:*TaEXPB5* instead of being fusiform in shape. The scale bars indicated are 0.5mm (A, B, C), 100μm (A1, B1, C1, A3, B3, C3), 50μm (A5, A6, B5, B6, C5, C6), and 10μm (A2, B2, C2).

### Slow degradation of the tapetum make KTM3315A male fertile at higher temperature

The tapetum is required for microspore development, and abnormal development of the tapetum can lead to pollen abortion and male sterility (Xie *et al*., 2014). To further understand whether inducing developmental defects of tapetum due to the silencing of *TaEXPB5*, affect pollen developing abnormally and male fertility in KTM3315A and make it not to occur fertility conversion at higher temperature, we observed anther tissue sections at five developmental stages for silencing plants and control plants by optical microscopy (Supplementary Fig. S4) and TEM (Fig. 7). At Tds, the microspore was enclosed in an aggregate with thick tapetum, and the middle layer were obvious and striped (Fig. 7A, F, K). In Euns, the middle layer became thinner due to degeneration, but it was still clearly visible (Fig. 7B, G, L). At the same time, the tapetum degraded obviously and the cell spacing became larger. When the tapetosome could be observed on the surface of the tapetum in silencing plants (Fig. 6B). With the development of microspore to Luns, the middle layer could hardly be seen in all materials (Fig. 6C, 6H, 6M). The tapetum degraded severely in silencing plants, and only a few prominent tapetum debris was remained (Fig. 7C), and tapetosome appeared on the surface (Fig. 7H, M). During Bns, the tapetum in silencing plants almost completely degraded and only some remaining tapetosome could be seen (Fig. 7D). While in control plants, there were still much and obvious tapetum. At Tns, only small and sparsely arranged tapetosome remained in silencing plants (Fig. 7E), but the large and densely arranged tapetosome remained in control plants (Fig. 7J, O). From the above results, we can speculate that slow degradation of tapetum may be the possible inducement of fertility conversion.

**Fig. 7.**
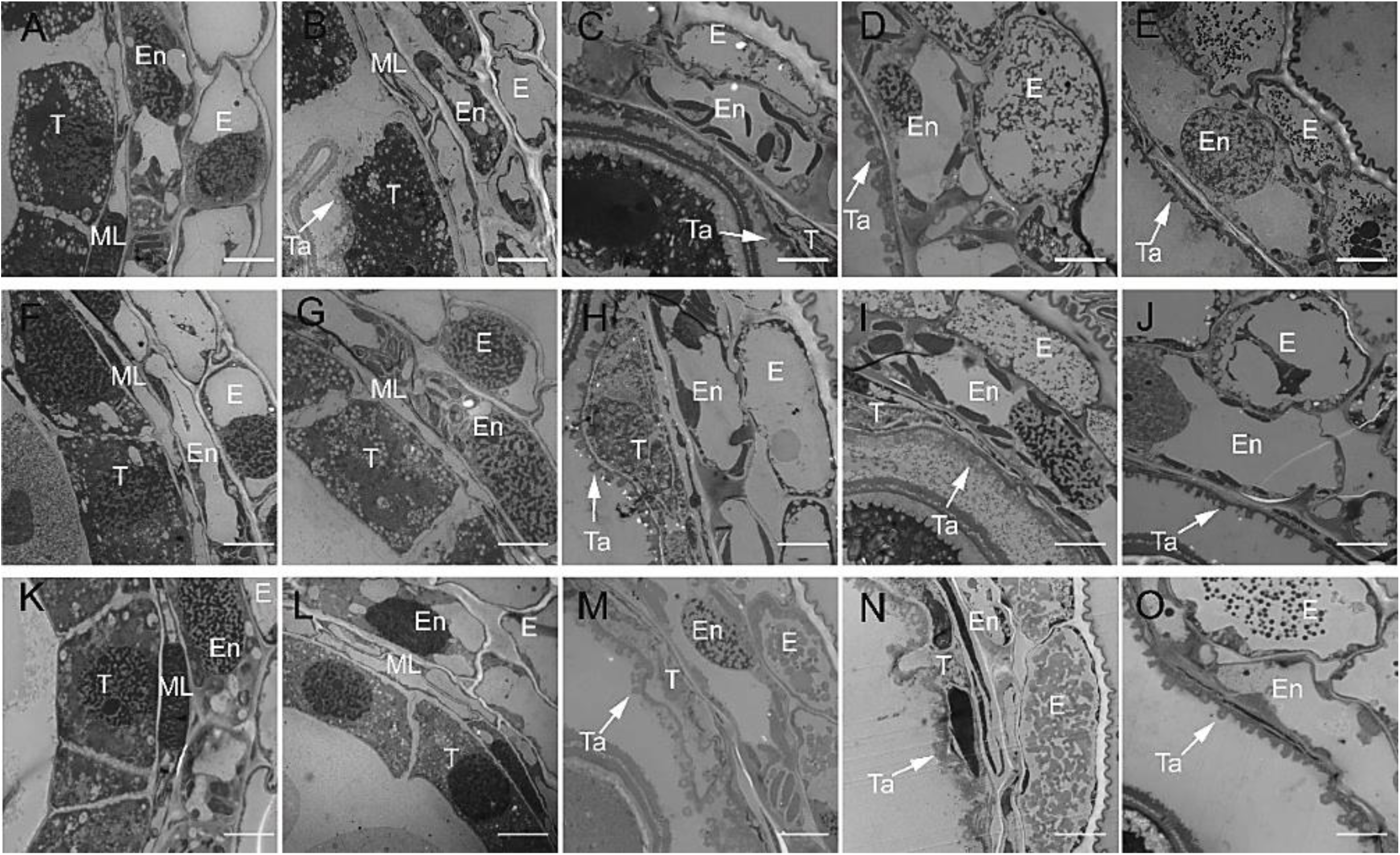
Comparisons of anther tapetum development by ultrathin section in silencing plants and control plants at different developmental stages. (A-E) silencing plants (BSMV:*TaEXPB5*); (F-J) control plants (AF); (K-O) control plants (BSMV:0); (A, F, K) Tds, tetrad stage; (B, G, L) Euns, early uninucleate stage; (C, H, M) Luns, late uninucleate stage, the tapetum degraded severely in silencing plants, and tapetosome appeared on the surface; (D, I, N) Bns, binucleate stage, in control plants, there were still much and obvious tapetum, only small and sparsely arranged tapetosome remained in silencing plants; (E, J, O) Tns, trinucleate stage, the large and densely arranged tapetosome remained in control plants. E, En, ML, T and Ta represent epidermis, endothecium, middle layer, tapetum and tapetosome, respectively. The scale bars are 2μm.

To further understand that the silencing of *TaEXPB5* leaded to abnormal degradation of tapetum, resulting in the loss of fertility conversion function under fertile conditions, we performed TUNEL assay to detect the degradation of tapetum in anthers. As shown in Figure 8, strong TUNEL-positive signals were detected in the tapetum of silencing plants at Tds (Fig. 8A), which means that the tapetum in silencing plants began to undergo apoptosis, while no apoptotic signals were detected in control plants (Fig. 8F, K). Until Euns, the tapetum of control plants began to show TUNEL green fluorescence signals (Fig. 8G, L), and as well as at the subsequent Luns, Bns and Tns, TUNEL apoptotic signals were detected in the tapetum of all anthers (Fig. 8C-E, H-J, M-O). The above result also was verified by DNA laddering assay (Supplementary Fig. S5). These results demonstrated that slow degradation of tapetum in KTM3315A under fertile condition was the potential inducement of male fertility.

**Fig. 8.**
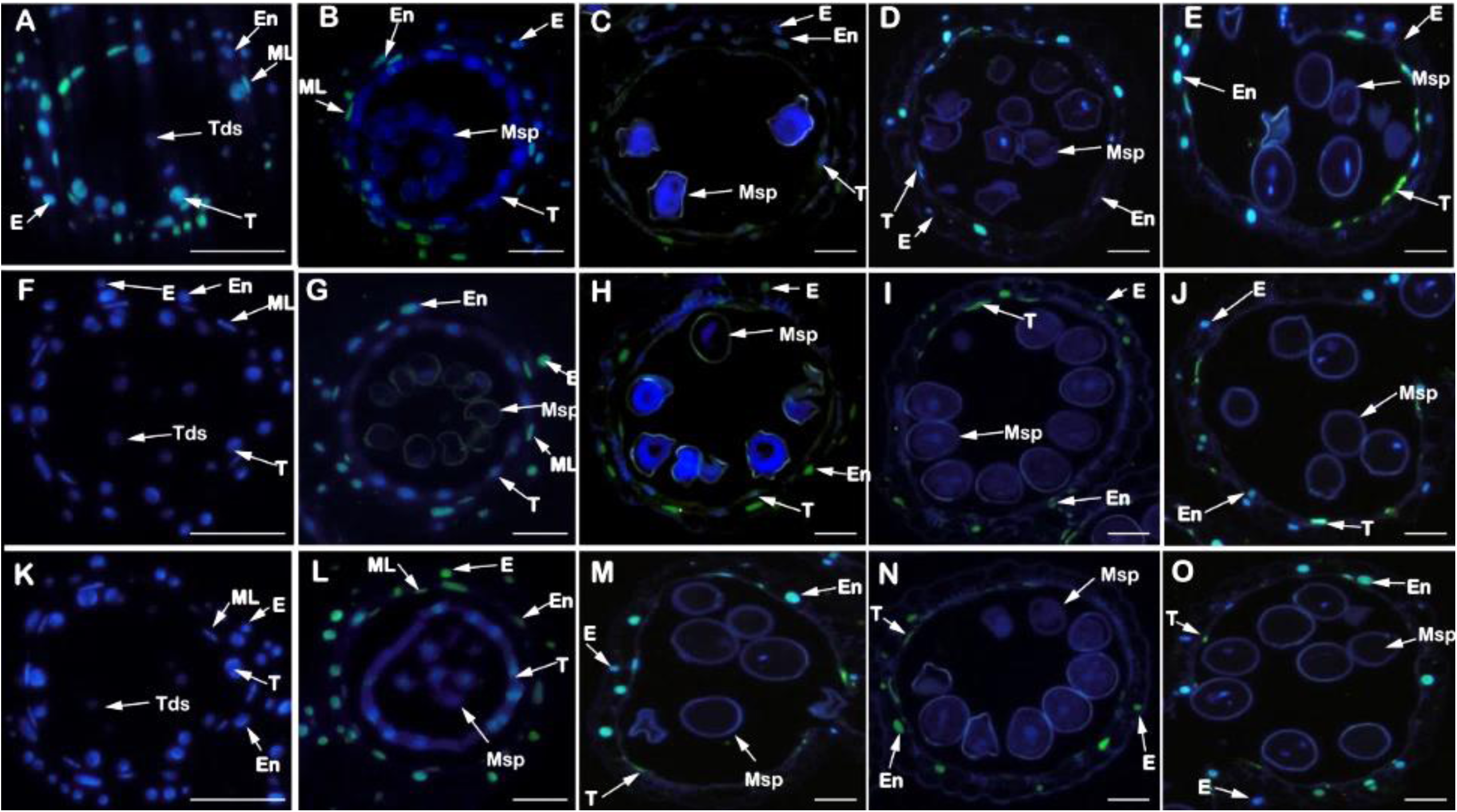
Detection of programmed cell death by TUNEL analysis in silencing plants and control plants at different developmental stages. (A-E) silencing plants (BSMV:*TaEXPB5*); (F-J) control plants (AF); (K-O) control plants (BSMV:0); (A, F, K) Tds, tetrad stage, strong TUNEL-positive signals were detected in the tapetum of silencing plants, while no apoptotic signals were detected in control plants; (B, G, L) Euns, early uninucleate stage, the tapetum of control plants began to show TUNEL green fluorescence signals; (C, H, M) Luns, late uninucleate stage; (D, I, N) Bns, binucleate stage; (E, J, O) Tns, trinucleate stage. E, En, ML, T and Ta represent epidermis, endothecium, middle layer, tapetum and tapetosome, respectively. The scale bars indicated are 50μm.

### The silencing of TaEXPB5 caused abnormal development of microspore exine, eliminating the effect of temperature on fertility under fertility conditions

Normal apoptosis of tapetum can ensure the normal development of microspore, and abnormal degradation of tapetum can cause male sterility (Falasca *et al*., 2013; Flores-Renteria *et al*., 2013; Papini *et al*., 2014). To determine whether the development of microspores was synchronized with the degradation of the tapetum in silencing plants and control plants, the epidermis of pollen grains at five developmental stages were observed by TEM (Fig. 9; Supplementary S6). At Tds, the development of microspores in all anthers were normal, and the primexine was just established (Fig. 9A; Supplementary S6A). One point of silencing plants differs from control plants was that the callose around the microspores has been degraded, while the callose in control plants were still well wrapped around microspores (Fig. 9F, K; Supplementary S6F, K). At Euns, there was obvious accumulation of sporopollenin on the surface of microspores. In silencing plants, sporopollenin was sparse and disordered on the surface of microspores (Fig. 9B; Supplementary S6B), while the accumulation of sporopollenin in control plants was normal (Fig. 9G, L, S6G, L). At Luns of microsporogenesis, the exine was formed, which was composed of nexine, baculum, tectum and sporopollen. There was still no obvious accumulation of sporopollenin in silencing plants, and the tectum was irregular (Fig. 9C; Supplementary S6C), indicating that sporopollenin accumulation was blocked and the structure of pollen wall was abnormal. On the contrary, many raised spores have accumulated on the surface of control palnts microspores, and the shape of the tectum was regular (Fig. 9H, M; Supplementary S6H, M). During Bns and Tns, the accumulation of sporopollenin, the morphology of the tectum, and the baculum in silencing plants were abnormal, and the entire exine was disorganized and thinner than that in control plants (Fig. 9D, E). The above results indicated that the silencing of *TaEXPB5* not only leads to the early degradation of tapetum, but also cause abnormal development of exine, which resulted in male sterility of silencing plants and loss of fertility conversion function.

**Fig. 9.**
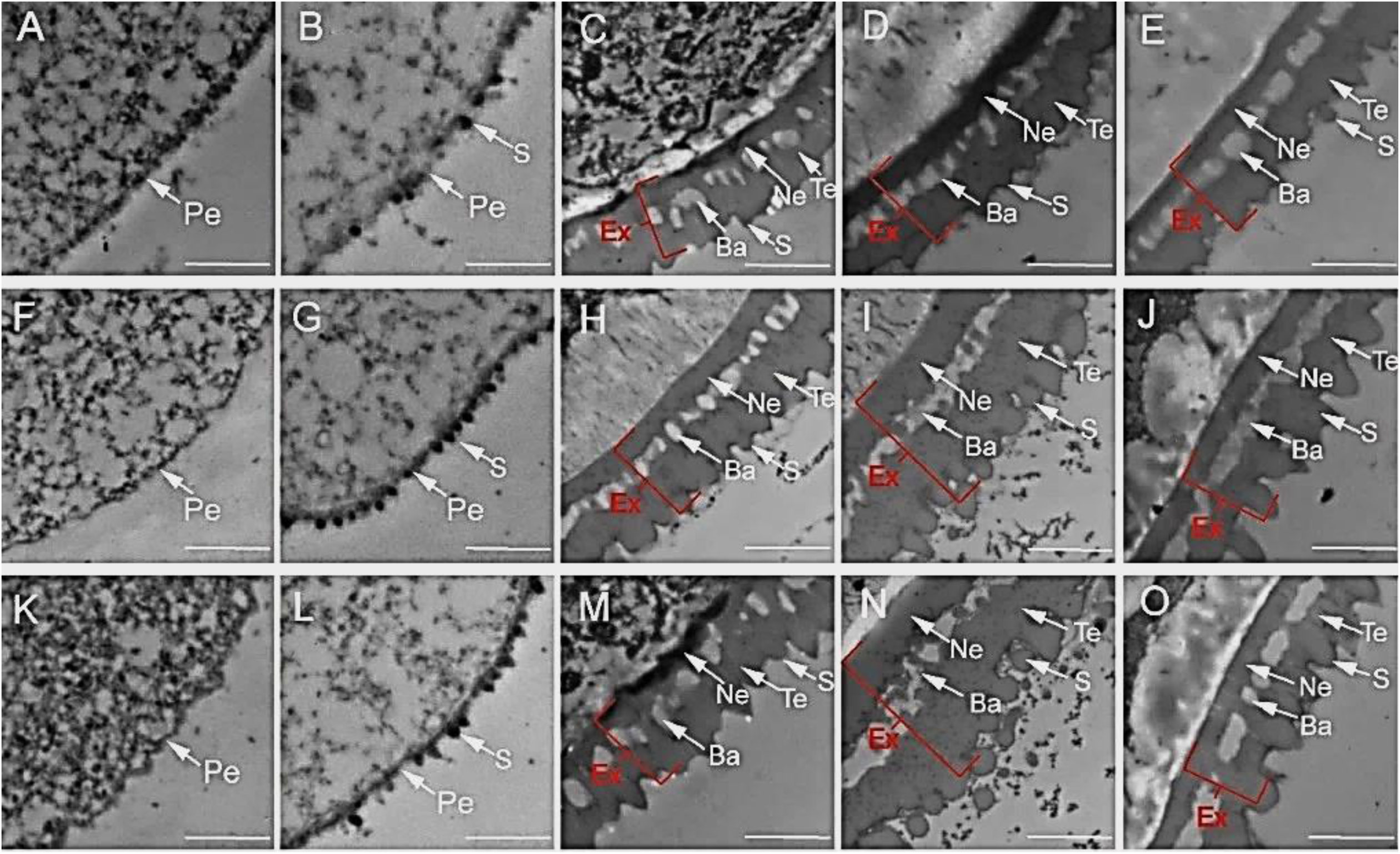
Development of pollen exine in silencing plants and control plants at different developmental stages. (A-E) silencing plants (BSMV:*TaEXPB5*); (F-J) control plants (AF); (K-O) control plants (BSMV:0); (A, F, K) Tds, tetrad stage, the callose around the microspores has been degraded, while the callose in control plants were still well wrapped around microspores; (B, G, L) Euns, early uninucleate stage, in silencing plants, sporopollenin was sparse and disordered on the surface of microspore, while the accumulation of sporopollenin in control plants was normal; (C, H, M) Luns, late uninucleate stage, there was still no obvious accumulation of sporopollenin in silencing plants, and the tectum was irregular, indicating that sporopollenin accumulation was blocked and the structure of pollen wall was abnormal; (D, I, N) Bns, binucleate stage, the entire exine was disorganized and thinner than that in control plants; (E, J, O) Tns, trinucleate stage. Ex, Ne, Ba, Te, Pe and S represent exine, nexine, baculum, tectum, primexine and tapetosome sporopollenin, respectively. The scale bars indicated are 2μm.

## Discussion

*TaEXPB5 conferred to pollen development and male fertility under fertile condition* The relaxation and expansion of cell walls are necessary for the growth of plant cells. Different from animal cells, plant cells are surrounded by cell walls, which are highly dynamic and complex networks (Vaahtera *et al*., 2019). Cells are always in a dynamic balance between the expansion of protoplasts and the restraint of cell walls, undergoing irreversible growth (Kuo *et al*., 2005). Expansion proteins are a type of cell wall binding protein that play an important role in the process of cell wall relaxation. Expansins usually have two typical domains, DPBB_1 and Pollen_allerg_1, and most of expansins have signal peptides (Liu *et al*. 2020). Similarly, TaEXPB5 contains both domains. Many studies have shown that the EXPA subfamily plays various roles in plant growth and development, such as *AtEXPA2* (Yan *et al*., 2014; Sánchez-Montesino *et al*., 2019; Xu *et al*., 2020), *AtEXPA5* (Son *et al*., 2012), *OsEXPA8* (Ma *et al*., 2013; Wang *et al*., 2014), *AtEXPA14* (Lee *et al*., 2013a), *AtEXPA17* (Lee *et al*., 2013b). However, researches on the roles of expansins in plant reproductive development are relatively limited. The expression loss of maize EXPB caused by transposon insertion can lead to pollen aggregation, resulting in poor powder dispersal when anthers were cracked, and pollen tube difficult to enter maize filament. This result was also verified in a transgenic maize plant that silencing all genes encoding EXPB subfamily extension proteins (Valdivia *et al*., 2009). In addition, the electronic expression profile analysis of *Arabidopsis* genes showed that the two expansin protein genes, *EXPA4* and *EXPB5* of the EXPB subfamily, were strongly expressed in dry pollen grains, aspirated pollen grains and pollen tube growth (Mollet *et al*., 2013). At the same time, in the detection of *AtAGP6*/*11* interactors based on expression and co-localization, it was found that *AtAGP6*/*11* can interact with other cell wall binding proteins, such as AtEXPB5, and participate in a variety of signaling pathways, in particular, endocytosis mediated plasma membrane remodeling pathway was involved in pollen development (Costa *et al*., 2013). In the GO analysis of tobacco and *Arabidopsis* expansin gene families, both *tobacco* and *Arabidopsis* EXPB subfamily were involved in sexual reproduction (GO: 0009553) (Ding *et al*., 2016). Previous studies have shown that AtEXPB5 has a two-stage expression pattern during pollen development. It is first expressed in pollen mother cells and tapetum before tetrad formation. Then the expression gradually decreased, and the signal disappeared after the microspores were released from the tetrad. In the second stage, the expression level suddenly increased at the mononuclear pollen stage and continued until the mature pollen stage (Lou *et al*., 2018). The expression pattern of *TaEXPB5* was similar to that of *AtEXPB5* in the second stage. Therefore, *TaEXPB5* conferred to pollen development and male fertility.

### TaEXPB5 is required for pollens development and male fertility under fertile condition

Plant male sterility is a phenomenon that functional pollen cannot be produced during staminogenesis due to physiological or genetic factors. There are various reasons for the formation of plant male sterility, such as disorder of sporogenous cell differentiation, microspore development and meiosis, and pollen or anther differentiation (Li *et al*., 2014). At present, the research on plant male sterility is mainly in cytology and molecular biology. The causes of sterility involve various aspects, including abnormal tapetum degradation (Ji *et al*., 2015; Yang *et al*., 2019), abnormal callose decomposition (Worrall *et al*., 1992), abnormal sporopollenin assembly (Li *et al*., 2020), abnormal pollen wall development (Yang *et al*., 2019), etc. In this study, we explored the causes of male sterility in Wheat by observing phenotypes and cytological sections. We found that the gene *TaEXPB5* silenced plants had withered anthers, shrunk microspores, advanced tapetum PCD, abnormal pollen outer wall formation and no anther dehiscence. Therefore, we can infer that the silencing of *TaEXPB5* eventually leads to the male sterility of KTM3315A under fertile conditions. In addition, the subcellular localization of *TaEXPB5* showed that it was expressed in cell membrane and nucleus. Tissue specific expression analysis showed that the expression of *TaEXPB5* was upregulated in the fertility anthers; In conclusion, according to the results of this experiment, we can conclude that *TaEXPB5* has important functions in anther development.

### Slow development of the tapetum make KTM3315A male fertile at higher temperature

The anther wall consists of epidermis, endothecium, middle layer and tapetum from outside to inside. Tapetum is the innermost cell of pollen sac wall, which is directly adjacent to sporogenous cells. It plays an important role in nutrition and regulation of the development and formation of pollen grains. Callose enzymes synthesized and secreted by tapetum can timely decompose the callose wall of tetrads, so as to separate mononuclear pollen grains from each other and ensure normal development. At the same time, it also has the function of synthesizing and secreting sporopollenin and transferring it to the outer wall of pollen grains (Papini *et al*. 1999). The related studies have also found that the abnormal PCD (program cell death) process of tapetum leads to the damage of pollen wall (Yi *et al*., 2016), and the early or delayed degradation of tapetum will cause male sterility (Li *et al*., 2006; Li *et al*., 2011). In addition, studies have also shown that the early degradation of callose can also lead to male sterility (Worrall *et al*., 1992). It is well known that male sterility is closely related to the early apoptosis of tapetum, as studied in rice (Ku *et al*., 2003; Li *et al*., 2004), sunflowers (Balk *et al*., 2001), maize (Rui *et al*., 2006). In addition, some genes specifically expressed in tapetum regulate tapetum development and affect anther development. For example, in petunia, the silencing of the gene *TAZ1* will lead to premature tapetum degradation and pollen abortion (Li *et al*., 2001); In *Arabidopsis, MS1* mutation leads to early tapetum degeneration and complete male sterility (Wilson *et al*. 2001). In rice, *PTC1* mutant showed delayed tapetum degeneration, abnormal pollen wall formation and termination of microspore development (Balk *et al*., 2001). In addition, barley *HvMS1* gene plays an important role in the development of anther tapetum. Silencing and overexpression of this gene will lead to male sterility in barley (Fernández Gómez *et al*., 2014). In our study, compared with the control, in the silencing plants of *TaEXPB5* gene, the first obvious phenotypic change was that the anther epidermis was shrinked and shriveled (Fig. 6A), and the surface of microspores was rough and shriveled (Fig. 6A3, A4). Through further cytological observation, it was found that the tapetum of silencing plants produced a strong apoptosis signal at the tetrad stage (Fig. 8A; Supplementary S5), and the degradation of tapetum was significantly earlier than that of control plants (Fig. 7, 8; Supplementary S4, 5). Therefore, we speculate that gene *TaEXPB5* causes delayed degradation of the tapetum, and resulting in male fertility, further demonstrated that *TaEXPB5* confer to male fertility conversion of KTM3315A under fertile condition.

### The silencing of TaEXPB5 causes abnormal development of pollen, which lead to loss fertility conversion function at higher temperatures

The plant pollen exine begins to form at the tetrad stage after meiosis. At this time, the primexine, which has been proved to act as a template during the construction of pollen exine, has just been produced between the microspore cell membrane and the callose wall (Heslop-Harrison, 2009). After the formation of primexine, sporopollenin will be orderly assembled and stacked under its guidance (Quilichini *et al*., 2014). *Male sterility 1* (*MS1*) is a plant homologous domain finger transcription factor, which has the function of regulating pollen primexine and tapetum programmed death in *Arabidopsis* (Ito et al., 2007; Yang et al., 2007). The *OSMS1* mutant of rice showed delayed degradation of tapetum and defects in pollen wall formation, and finally manifested as male sterility (Yang *et al*., 2019). The Poaceae specific gene *EPAD1* mutation in rice destroys the continuity and homogeneity of the primexine layer at the tetrad stage and the arrangement of the primary columnar layer, thus changing the assembly pattern of sporopollenin on the pollen surface and eventually leading to pollen abortion (Li *et al*., 2020). In this study, the silencing of *TaEXPB5* destroyed the continuity and homogeneity of the sporopollenin arrangement on the pollen surface of the late uninucleate stage. In the next development process, the abnormal structure of pollen wall showed that the surface was disordered and not smooth, the tectum was irregular, and the assembly pattern of sporopollenin changed. Finally, resulting in male sterility. Therefore, we speculate that the gene *TaEXPB5* is indispensable for the fertility of thermo-sensitive male sterile wheat line KTM3315A.

### BSMV-VIGS can effectively verify the function of wheat genes

Tissue culture and regeneration remain rate-limiting steps in plant gene editing. For some monocots, especially common wheat containing a large and complex hexaploid genome, gene editing is particularly difficult (Ran *et al*. 2017; Gao 2021). However, virus-induced gene silencing (VIGS) has been developed as a reverse genetic tool for rapid identification of plant gene functions in recent years. Compared with traditional transcription-level gene silencing technology, VIGS technology has many advantages, such as fast, effective, simple operation and wide application range (Zhang *et al*., 2016). BSMV was selected as a vector for plant genetic engineering due to its wide host range, seed inheritance and high replication rate in vivo. Mechanical damage is the main method of BSMV infection (Lee and Kanyuka 2012). The virus is named for the irregular dark green and light green stripes produced when it infects barley. In the early stage of infection, yellow or white fading green stripes appear at the base of new leaves, and then gradually spread to the whole plant, accompanied by dwarfing phenotypes. So far, BSMV-VIGS has identified many genes related to stress resistance and disease resistance, such as *NtMYB3* (Ping et al. 2021), *CYP96B22* (Delventhal *et al*., 2014), *TaADF7* (Fu *et al*., 2014), *ScBx1* (Groszyk *et al*., 2014), *TaEIL1*(Duan *et al*., 2013), *TaBTF3* (Kang *et al*., 2013). Therefore, VIGS technology was used in this study to study the function of the gene *TaEXPB5* related to pollen development, and the results showed that *TaEXPB5* playing an important role in the process of pollen fertility conversion. The BSMV-VIGS system down-regulated the KTM3315A pollen development-related gene *TaEXPB5*, which reduced its expression in anthers and failed to complete pollination, leading to decreased fertility. This research lays the foundation for the genetic improvement of future crops.

## Conclusion

In this study, the quantitative analysis assay of gene *TaEXPB5* screened by studying the transcriptome and proteomics of temperature sensitive male wheat KTM3315A at different fertility temperatures showed that which positively regulated the fertility of KTM3315A. Cytological observation of anthers in five stages of silencing plants and control plants showed that, after gene silencing, which affects the development of microspore and the fertility of pollen by regulating the early degradation of anther tapetum and the formation of sporopollenin. Therefore, the genes *TaEXPB5* have an important role in anther development. The silencing of the gene can lead to a decline of fertility. The data provide new insights into the mechanism involved in wheat pollen development for thermo-sensitive male sterility.

## Supplementary data

**Table S1** The primer lists in this study.

**Fig. S1** Sequence of *TaEXPB5* in KTM3315A under fertility conditions.

**Fig. S2** Amplification of *TaEXPB5* silencing fraction and the result of transcription in vitro for BSMV vectors and recombination vectors.

**Fig. S3** Relative expression in silencing plants and control plants.

**Fig. S4** Comparisons of anthers and tapetum in silencing plants and control plants at different developmental stages.

**Fig. S5** Detection of programmed cell death by DNA laddering analysis in silencing plants and control plants at different developmental stages.

**Fig. S6** Development of pollen exine in silencing plants and control plants at different developmental stages.

## Acknowledgements

This work was supported by the National Natural Science Foundation of China (31771874, 32072060). We thank Beijing Biomarker Biotechnology Co., Ltd (China) for their assistance with data processing and bioinformatics analysis, PTM BioLab, Inc. (China) for their assistance with data processing.

## Author contributions

XS and LZ contributed to the study conception and design. Material preparation, data collection and analysis were performed by XG, XW, XY, and JW. The first draft of the manuscript was written by XG and XW, all authors commented on previous versions of the manuscript. All authors have read and approved the manuscript.

## Data availability

The data supporting the findings of this study are available within the paper and within its supplementary materials published online.

